# Assembling bacterial puzzles: piecing together functions into microbial pathways

**DOI:** 10.1101/2024.03.27.587058

**Authors:** Henri Chung, Iddo Friedberg, Yana Bromberg

## Abstract

Functional metagenomics enables the study of unexplored bacterial diversity, gene families, and pathways essential to microbial communities. However, discovering biological insights with these data is impeded by the scarcity of quality annotations. Here, we use a co-occurrence-based analysis of predicted microbial protein functions to uncover pathways in genomic and metagenomic biological systems. Our approach, based on phylogenetic profiles, improves the identification of functional relationships, or participation in the same biochemical pathway, between enzymes over a comparable homology-based approach. We optimized the design of our profiles to identify potential pathways using minimal data, clustered functionally related enzyme pairs into multi-enzymatic pathways, and evaluated our predictions against reference pathways in KEGG. We then demonstrated a novel extension of this approach to predict inter-bacterial protein interactions amongst members of a marine microbiome. Most significantly, we show our method predicts emergent biochemical pathways between known and unknown functions. Thus, our work establishes a basis for identifying the potential functional capacities of the entire metagenome, capturing previously unknown and abstract functions into discrete putative pathways.

## 1 Introduction

The rise in high-throughput sequencing technologies has revolutionized the study of microbial communities via a marked increase in the quantity and quality of microbial sequence information [1, 2]. However, the vast majority of microbial life remains uncultured and undescribed. This unknown microbial world represents, by some estimates, up to 87% of bacterial genera on Earth [3, 4]. The sheer abundance of corresponding bacterial DNA and protein sequences invites new types of analyses to identify unknown microbial functions and their interactions. While significant efforts have been made to characterize the functions represented in this space [5–9], there have been few attempts at connecting these functions as interacting groups, or pathways [10, 11].

Pathways, comprising proteins that participate in interconnected biochemical reactions, play pivotal roles in the metabolic activities essential to life. Bacterial pathways underpin fundamental processes necessary for all life on Earth: the synthesis of cellular components, environmental nutrient cycling, and energy flow. Understanding these complex pathways requires assessment not only of the known microbial proteins, which represent a minority of total bacterial function, but also consideration of the myriad unknown functions yet to be discovered [12–15]. The challenge, therefore, lies in identifying the interactions of bacterial proteins without requiring prior knowledge of their functionality. Co-occurrence analyses — the counting of paired instances within a collected sample — have long been used to understand the relationships between objects, without explicit awareness of their individual roles [16–22]. Here, we used the co-occurrence of putative microbial functions, both known and unknown, to (1) identify networks of proteins that recapitulate pathways endowed by bacterial genomes and (2) predict emergent microbiome functionality from metagenomic data.

Earlier studies have used patterns of gene co-occurrence within an organism to predict putative functional interactions [23, 24]. They quantified co-occurrence of genes by comparing gene phylogenetic profiles, i.e. the presence or absence of specific genes across different organisms. By analyzing the evolutionary relationships and patterns of gene distributions in this manner, these studies identified genes that may be functionally related or involved in similar biological processes across species. Phylogenetic profiles are *N* -length vectors of orthologous genes that are present or absent across *N* species. The similarity of two such profiles can be used to quantify the cooccurrence of any two genes and to label pairs of genes as functionally interacting. This approach reasons that functionally linked genes co-evolve, and therefore, will co-appear in the same subset of organisms.

Gene phylogenetic profiles have been successfully used to infer pairwise protein interactions and to, subsequently, inform multi-protein interaction networks, infer evolutionary relationships, and label protein subcellular localization [25–30]. Studies have evaluated the impact of profile length, organism selection, and similarity metrics as adjustable parameters of profile construction [31–34]. Refinements of phylogenetic profiling have explored the additional integration of gene expression values, accounting for the random probability of co-occurrence, as well as non-homology-based profile element identification [29, 35, 36]. Note that building phylogenetic profiles for a gene or a protein requires the identification of its orthologs. In the original study, orthologs for a protein of interest were identified as those matching the query sequence with a score above an alignment threshold relative to the size of the searched database [24]. Since then, profile elements have been identified using bit-score thresholds, protein domains, membership in Clusters of Orthologous Groups of proteins (COGS), and methods for distinguishing between orthologs and paralogs [25, 34, 37–40].

Here, we constructed profiles using functionally-similar proteins, regardless of their orthology status; as described in Mahlich *et al*. [41]. In previous work, we developed the FUnctional-repertoire SImilarity-based Organism Network or *fusion* — a clustering of proteins by functional similarity, which improved labeling of function compared to other sequence similarity-based methods [41, 42]. We assumed proteins of one *fusion* cluster to be of the same function, and these were then used to build profiles in a manner similar to orthologs in previously explored phylogenetic profiles. We compared our *fusion* results to those of profiles that use alignment-based ortholog detection at varying sequence similarity thresholds. We found that using *fusion* improved prediction accuracy over alignment-based profiles for a labeled set of enzymes and corresponding pathways.

Importantly, *fusion* clusters allow for the identification of functionally similar proteins without pre-existing functional annotations. This results in a prediction-discovery process that provides testable hypotheses for experimental validation.

We further optimized our previously developed *faser* method for labeling sequencing reads to identify *fusion* functions encoded in metagenome data[43]. We were therefore, able to build functional profiles of microbiomes, allowing for the discovery of yet-unseen, emergent, microbiome-specific pathways. When applied to marine microbial metagenomes, our approach recognized energy metabolism-relevant interactions between proteins from different organisms. That is, *fusion*-based profiles were able to identify pathways specific to individual bacteria and, unlike other methods, pick out pathways shared between organisms. These latter pathways are likely characteristic of the unique functional capabilities of the microbial communities specific to a given environment, addressing the fundamental microbiome question “What are these genes doing?” Using functional co-occurrence to identify metagenome-encoded pathways can help shed light on the metabolic capabilities of microbial communities, enabling a more comprehensive understanding of the microbial world.

## 2 Methods

### 2.1 Workflow

### 2.2 Fusion data set

As stated, this work builds upon our previous study, which established and validated our unique bacterial protein function annotation scheme — *fusion* [41]. Here, we used a subset of the Balanced Organism data set, assembled as a concise and representative collection of bacterial functions. Briefly, we retrieved the proteomes of 8,906 distinct bacterial assemblies from NCBI GenBank, comprising *∼*31.5 million proteins. From this set, we removed redundant identical sequences and sequences shorter than 23 amino acids (the limit for identifying protein functional similarities [44]). We grouped the remaining *∼*15.5 million proteins into *fusion* clusters using a multi-level clustering method based primarily on the proteins’ pairwise Homology-derived Functional Similarity of Proteins (HFSP) score [44]. The resulting clusters, referred to as *fusion* clusters, are groups of proteins with similar functional capabilities. To avoid overestimating protein co-occurrence due to over-representation of closely related species in our data, we further reduced this set, using Treemmer [45], to a taxonomically balanced set of 1,502 distinct organism genome assemblies that retain at least 90% of total genetic diversity of the full set.

Note that at the time of the analysis reported here, 109 of 1,502 organism assemblies present in the original balanced data set had been removed or suppressed in NCBI databases, and, therefore, were not considered here. To reduce the influence of proteins specific to individual bacteria, we filtered the Balanced Organism set to retain proteins found in a least five of the remaining 1,393 organisms. The final Balanced *Fusion* data set used in this work consisted of 1,393 assemblies and 3,613,779 proteins grouped into 77,042 *fusion* clusters (Ref Tab. S1).

### 2.3 *fusion* Profiles

We made and used *fusion* profiles to predict and quantify functional interactions between proteins based on their co-occurrence across organisms. We constructed these profiles by noting the presence/absence of proteins belonging to the same *fusion* cluster across bacterial genomes in our data set. Thus, each profile is a *fusion* cluster-specific *N* -element binary vector, where *N* is the number of bacterial genomes (1,393), with 1 or 0 entries indicating the presence or absence, respectively, within each bacterium of a protein from this cluster. Note that each protein in a particular *fusion* cluster has the same fusion profile.

We retained 306,377 protein-profiles for the enzymes in the Balanced Organism Data set that had experimentally-derived EC number annotations (resolved to all 4 EC numbers, extracted from Swiss-Prot [46, 47]), or that shared an HFSP score *geq* 20 with one of these proteins; the considered ECs had to participate in a KEGG module. Note that we had previously found that an HFSP score *geq* 20 accurately identified shared enzymatic function with 95% precision [44]. These proteins had 1,717 unique EC numbers, distributed across 1,661 *fusion* clusters. Proteins of 401 EC numbers were found in more than one *fusion* cluster and 345 clusters contained more than one EC number.

### 2.4 KEGG Reference Pathways

Here we used KEGG as the ground truth reference set of bacterial pathways; we therefore retrieved a list of complete KEGG modules for each organism in our data set. A KEGG module is a set of enzymes (with associated EC numbers) that participate in a specific metabolic pathway. Pairs of proteins were assigned positive and negative labels if they participated in the same module in a given organism, as was done in previous assessments of phylogenetic profile performance using KEGG modules [48–50].

Based on module participation, we calculated positive-labeled data as the total number of unique protein pairs taken from the set of proteins *within each module, P*_*ij*_, with *i* ranging from 1 to *N* organisms and *j* ranging from 1 to *M*_*i*_, or the number of Modules in organism *i* (Eqn. 1). We then calculated Negatives by compiling all unique possible protein pairs *between different modules* within the same organism. Here, *j* and *k* both index *M*_*i*_ modules, except when *j* = *k*, and *i* ranges from 1 to *N* total organisms. In the Negatives, each protein, *p*_1_ or *p*_2_, in the pair is from a different module (Eqn. 2).

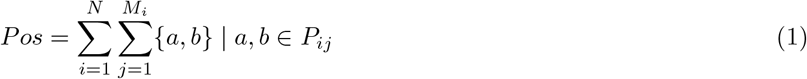

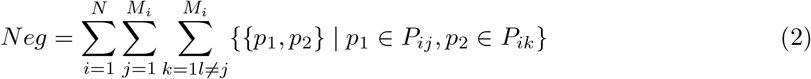

For each module, we also retrieved the EC numbers of component proteins, their KEGG Ortholog (KO) assignment, and the number of reaction steps in the module [51]. A KO is an orthologous group of genes that are assumed to perform a specific molecular function or participate in a particular biological process across different species. Thus, a KO assignment is an additional reference for the specific function of a given protein. For each module, we identified pairs of proteins that catalyze the same module step but belong to different KOs. We labeled pairs of these functional analogs as non-orthologous replacements or NORs.

### 2.5 Evaluating Profiles

To assess any improvement generated by using *fusion* for profile element detection over sequence similarity in phylogenetic profiles, we compared the protein interaction predictions of *fusion* profiles to those of profiles made with MMseqs2, a software suite for searching and clustering large protein databases [2]. We used MMseqs2 to identify orthologous proteins by their alignment bit scores. Parameters for MMseqs2 search were selected to match PSI-Blast sensitivity. We used a *k*-mer (`-k`) size of 7, *k*-mer threshold (`--k-score`) of 85, limited search to E-values below 10^*−*5^, sensitivity parameter (`-s`) of 7.5, and max sequence return (`--max-seqs`) of 4,000. We then applied different bit-score thresholds (40, 60, 80, and 100) to determine orthologous proteins. The presence of one or more matches above the threshold in an organism’s genome is 1 in the profile and 0 otherwise. We evaluated each method’s performance by its ability to identify reference bacterial protein interactions based on profile similarity. The reference interactions were defined as proteins participating in the same KEGG module as described above.

#### Evaluating profiles

To evaluate whether either method’s prediction performance correlated with the accuracy of identifying pairs of functionally similar proteins, we calculated the Rand Index for *fusion* clusters and MMseqs2 annotations, comparing each to multiple definitions of function. The Rand Index is a count of the number of protein pairs that share the same label, *a*, over the total number of possible protein pairs in set *n* (Eqn. 3) [52]. For both profile-building methods, we determined the number of protein pairs, that (1) share the same EC number at three levels, (2) share the same EC number at four levels, (3) share the same KO label, and (4) participate in the same KEGG module, and (5) are NORs for the same reaction step in a module. Protein pairs where one or both of the proteins did not have a corresponding function label were not considered. To quantify the influence of sequence similarity, we also measured the average sequence identity of all pairs of proteins; we used the MMseq2 align command with `--alignment-mode 3` to measure the number of identical aligned residues divided by the total (gapped) alignment length.

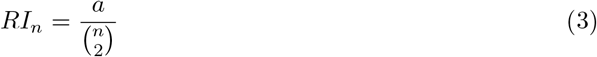

#### Evaluating predicted pairwise interactions

We first evaluated predictions as binary classification tasks for all pairs of proteins, to answer the question: are these two proteins included in the same KEGG module? To this end, we measured the Jaccard similarity between any two proteins’ profiles (Eqn. 4), predicting pairs whose Jaccard similarity was above a specific threshold as participating in the same module.

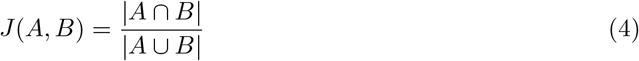

We assessed our predictions using measures of precision, sensitivity, specificity, and *F*_1_ (Eqns. 5–8). As described above, all reference protein pair interactions (Positives) were evaluated per organism, cumulatively summing to the total number of positives for method evaluation. Note that, this approach to generating the interacting protein pair ground truth (as described above), while directly comparable to prior functional interaction studies using phylogenetic profiles [48, 50, 53– 55], skews the number of putative positive and negative interactions. First, because KEGG modules are discrete functional units of larger pathways, this approach discounts the possible interactions between proteins in different modules of the same pathway, overestimating the total number of negatives. At the same time, as we consider any pair of proteins in a module to be interacting, the number of incorrectly labeled positives increases non-linearly with the number of proteins in a module. This method of labeling interacting pairs negatively biases the estimates of any prediction method’s precision and, arguably significantly more so (due to module sizes), recall.

We therefore propose an additional performance measure, *Group Recall*, which reflects the number of unique proteins within a module, instead of pairs of proteins. Group Recall labels as Positives the number of unique, module-specific proteins predicted to interact with other module proteins. We compare Pair Recall and Group Recall to estimate the effect of the pair-based evaluation bias.

Further, to accommodate our incomplete knowledge of bacterial life on Earth, we looked to determine the performance of *fusion* and MMseqs2 profile-based predictions in the presence of different numbers of available organisms. We compared the scores of the 1,393-size profile against the scores of smaller profiles of sizes 50, 100, 300, 500, 700, 900, and 1100. When evaluating smaller profiles, we created 100 replicates of each profile length below 1,393 by randomly selecting a subset of the total organisms. Since each profile length replicate uses a unique set of organisms, the number and position of filled elements for a protein’s profile varies between replicates.

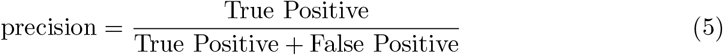

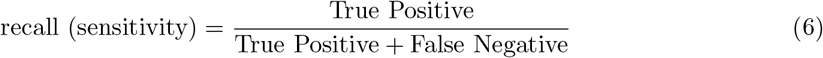

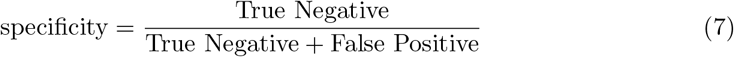

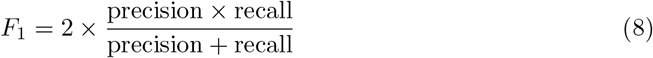

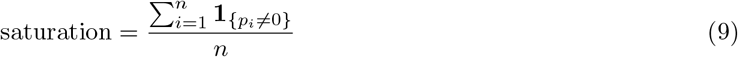

Ubiquitous or extremely common proteins found in most bacterial assemblies will, not unexpectedly, have high Jaccard similarities to other commonly found proteins, even if they do not participate in the same pathway. To reduce the number of these false positives, we defined a saturation filter for each protein that measures the percentage of filled elements in their profiles. We then applied this filter to predicted protein pairs, and if one or both of the proteins exceeded the saturation filter value (Eqn. 9), the prediction was removed from the evaluation. Note that the saturation level of proteins is a fraction. The numerator is the sum of *n* elements in profile *p*, and each element *i* has a value of 1 or 0. The denominator is the total number of elements *n*. Therefore, the number of proteins removed between profiles of different lengths and replicates of the same length varied. We tested saturation thresholds from 0.5 to 1 in increments of 0.1 to determine the optimal saturation threshold for accurate prediction.

#### Evaluating complete pathway predictions

We generated and clustered networks of interacting proteins using both *fusion* and MMseqs2 profiles. Here, proteins were nodes and the Jaccard similarity between the corresponding profiles was represented as the weighted edge between nodes. We converted each edge in our network into a pair of directed edges and clustered the resulting networks using HipMCL (High-performance Markov Clustering), a distributed-memory algorithm implementation of Markov Clustering, building a different network per organism and labeling the resulting clusters as potential pathways [56, 57]. We used the default parameter values for selection number (S = 1,100) and recovery number (R = 1,400). The S and R parameters prevent the pruned matrix during MCL from becoming too dense or sparse and indicate the number of column elements to prune or retain respectively. We varied the inflation parameter *I* from 1–4 to determine optimal performance, and identified an inflation parameter of 4 as best for our application.

To evaluate how well each set of predicted pathways recapitulated KEGG modules, we scored predicted putative pathways by identifying the pathways with the highest Jaccard similarity to a given KEGG module. For evaluating pathway prediction in individual organisms, we considered proteins in a network cluster from the same organism as interacting, regardless of EC annotation. For assessing emergent microbiome pathways we considered all proteins and, thus, possible pathways together. For the latter process, we excluded from consideration proteins that co-occur in a known KEGG pathway.

### 2.6 Metagenome data and short read assignment

We hypothesized the co-occurrence of functions across different microenvironments could be used to identify pathway interactions in the microbiome, in the same manner as phylogenetic profiles have been shown to identify pathways in individual organisms (Ref Fig. 2). We therefore extended our application of *fusion* profiles to identify emergent pathways in microbial communities. We compared the co-occurrence of functions across 480 marine microbial metagenomes from the bio-GEOTRACES project (ProjectID: PRJNA385854, https://www.ebi.ac.uk/ena/browser/view/SRX2974736) [58]. These samples came from 91 different locations and were collected in depths of 2–10 meters. Metagenomic data was generated using Illumina NextEra XT kits and sequenced on an Illumina NextSeq 550. The complete data set contains over 5 terabases encompassing 1.67 *×* 10^10^ paired-end reads of raw sequence data.

**Figure 1.**
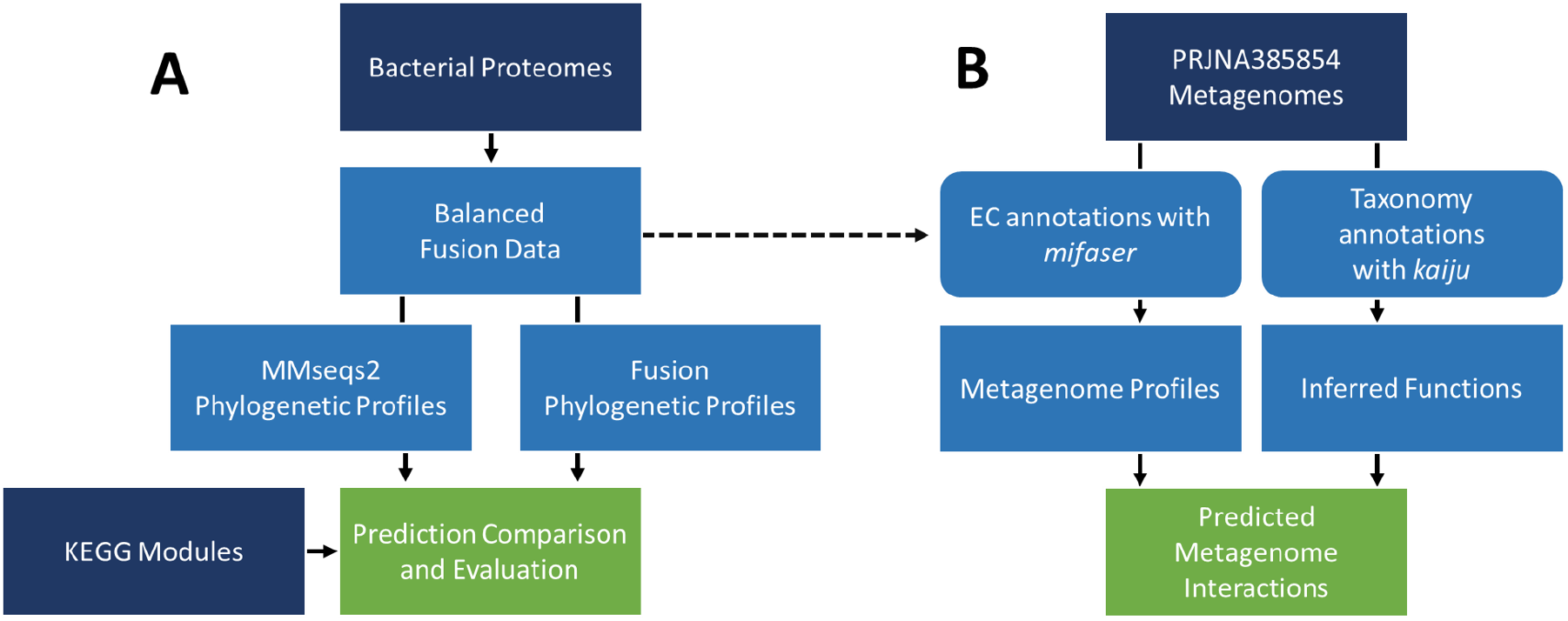
Project Workflow. A) Phylogenetic profiles were built from a taxonomically balanced set of bacterial proteomes using two methods: *fusion* and MMseqs2. We estimated the Jaccard profile similarity threshold for predicting protein interactions from these data. The predicted interactions were compared against KEGG modules, and evaluated via their precision, recall, and *F*_1_ score. B) We applied the same method for profile comparison to metagenome data. We annotated metagenomic data using two methods. (1) We used the *faser* method to map metagenomic reads to corresponding *fusion* clusters that contain proteins with annotated EC numbers. (2) We also used the Kaiju method to first annotate metagenome taxonomic composition and inferred present ECs from the identified taxa via KEGG. In both cases, functions with similar metagenomic profiles that did not co-occur in a single organism were predicted as potential novel interactions.

**Figure 2.**
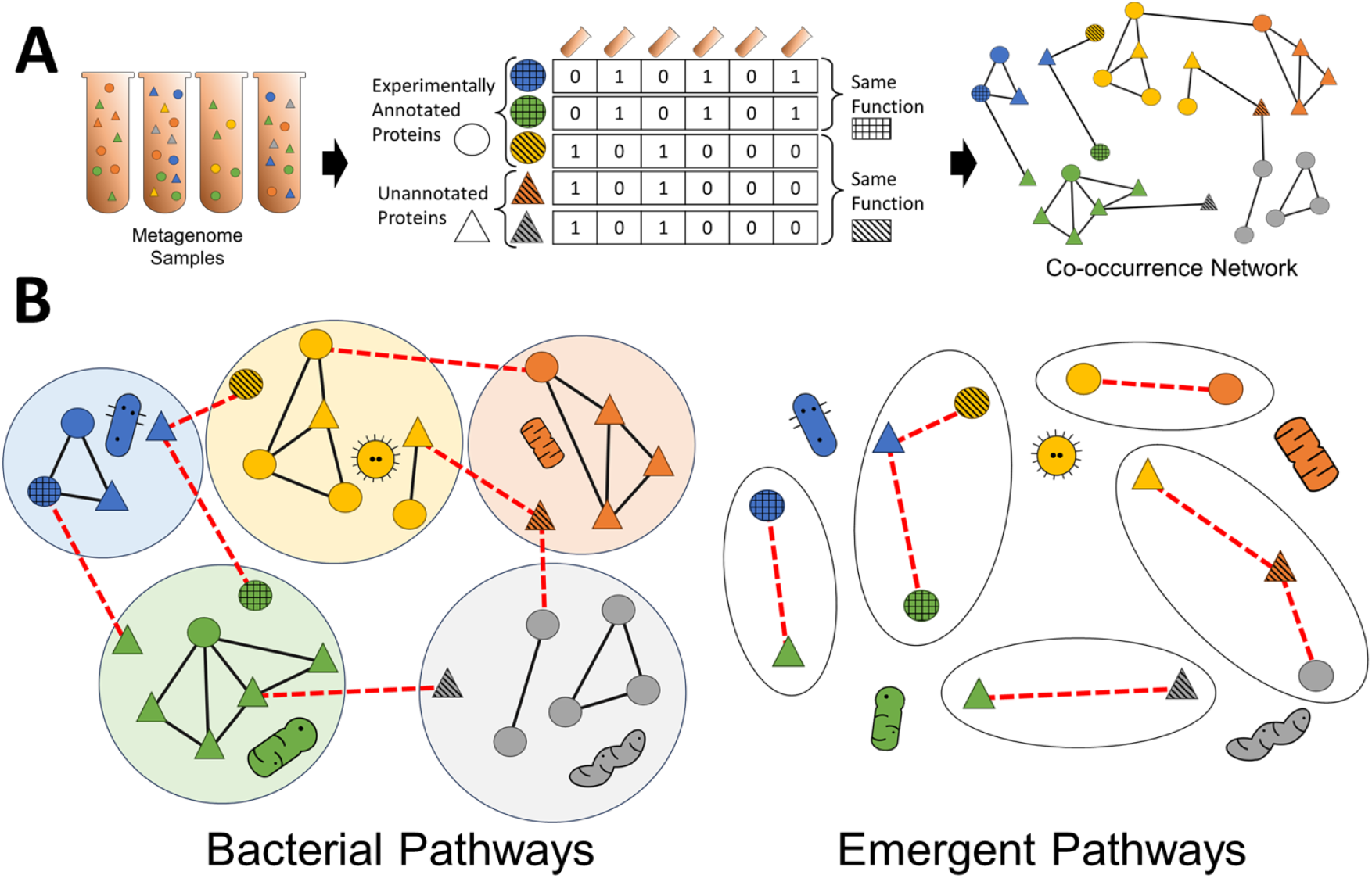
Metagenome Pathway Detection Approach. A) Using *faser* to parse metagenomic reads, we labeled putative *fusion* clusters, or functions, present in microbiome samples. We then created metagenome profiles, counting the co-occurrence of *fusion* clusters across different samples. Cross-hatching indicates whether proteins share the same *fusion* cluster, or function, and proteins in the same cluster will necessarily have identical profiles. Circles and triangles are used to distinguish between experimentally annotated and unannotated proteins. Color is used to represent proteins originating from different bacteria. We cluster metagenome profiles using their Jaccard similarity as edge weights. B) Identifying bacterial and emergent pathways based on identifying clusters of proteins within in the same organisms or exclusively between organisms, respectively.

For these samples, we created *metagenomic profiles*, representing the presence or absence of functions across different microenvironments. We then calculated the Jaccard similarity between any two functions’ metagenomic profiles to infer functional relationships.

Essential to our profile creation approach is the direct mapping of reads to function, without prior assembly or assignment to known taxonomy. To achieve this, we used *faser*, a fast, high-precision method for annotation of molecular functionality citeZhu2017faser. With *faser*, we mapped the metagenomic reads to functional annotations — both to EC numbers and *fusion* clusters [42]. The *faser* pipeline translates and aligns input read sequences to a reference database of proteins of annotated function. In its original application, *faser* was tested for accuracy of assignment of EC numbers. For use in our workflow, we evaluated *faser* performance in assigning reads to *fusion* clusters.

We split the *fusion* data set by organism (1,393) into 70:30 training (989) and testing sets (404). For each organism in the test set, we split their largest complete genomes into 10,000 random fragments of lengths 50-250 base pairs (4,040,000 fragments), representing possible genomic reads. Fragments not in coding regions (8% of total fragments) were then removed from consideration. We then used *faser* to label fragments in the coding regions with *fusion* cluster IDs of the training set proteins. We evaluated *faser*’s performance at correctly labeling test reads’ *fusion* clusters using the default *faser* performance cutoff of 20. The *faser* pipeline precisely annotated the fragments, achieving a precision of 0.94, recall of 0.52, and an *F*_1_ score of 0.67 – a performance similar to that of the originally reported values [43].

Following our evaluation, we used all proteins from the Balanced *Fusion* set as the reference database for *faser* metagenome analysis. Note that our dataset replaced the original smaller experimentally annotated database of the *faser* method [43], allowing us to capture a wider set of functionality.

In analyzing metagenomes, our goal was to predict functional relationships between proteins from different organisms, i.e. emergent functionality, based on their co-occurrence across multiple metagenomes. We hypothesized these emergent pathways to be putative ecological interactions between microbes, where a metabolite, or the protein itself, produced by one microbe is made available to other microbes in a multi-organism interaction.

To focus on such putative emergent functions, for all predictions of interactions, we filtered out pairwise interactions known to occur in individual organisms present in our metagenomes. For this, we used Kaiju [59], a fast and sensitive metagenome taxonomic classification program, to assign each read to a taxon in the NCBI taxonomy. For the reference database, we used the RefSeq data set provided by the developers of Kaiju (July 3, 2022), consisting of protein sequences from genome assemblies of Archaea and bacteria with a Complete Genome assembly level, as well as viral protein sequences from NCBI RefSeq. To ensure confidence in the presence of detected taxa, we filtered annotations to taxa that comprise *≥* 1% of annotated reads per file. We then disregarded any functional relationships from proteins found in pathways present in these taxa.

As described above for phylogenetic profiles, we calculated the Jaccard similarity between metagenomic profiles of EC functions identified with *faser*. We only considered profiles for functions found in at least two metagenome samples to avoid counting overly specific or unique functions. We compared Jaccard similarities for metagenome profiles and predicted them as interacting pairs of functions when similarity was *≥* 90%. We then generated networks from the metagenome profiles in the same manner as *fusion* and MMseqs2 profiles: converting the similarities between profiles into pairs of directed edges and clustering the resulting networks with HipMCL. We also generated randomized comparisons of our networks by permuting the edge weights between nodes before clustering. We then evaluated how well the resulting clusters encapsulated expected KEGG modules from known present bacteria, and if they indicated the presence of modules from undetected species or potential inter-species interactions.

## 3 Results

### 3.1 Profile parameter optimization improves prediction of functional interactions

We first evaluated how well our pathway predictions, made by *fusion* and MMseqs2 profiles, reflect our gold standard pairwise interactions extracted from KEGG pathways (Methods). We calculated the highest *F*_1_ score for each prediction-threshold/method/parameter-combination, reporting the mean over 100 replicates when comparing profiles of lengths less than 1,393. When *F*_1_ was comparable between different profiles (*±*0.01), we prioritized precision as a performance measure to give a higher confidence in downstream analyses.

Specifically, we optimized predictions by varying parameters of profile length and saturation. Profile length refers to the total number of organisms used to create the profile, and saturation reflects the fraction of organisms that carry out a particular function. Exploring the profile length allowed us to estimate the impact that our limited knowledge of existing organisms had on pathway identification. The performance of both *fusion* and MMseqs2 profile-based methods have improved for longer profiles (Ref Fig. 3D-F), i.e. both pathway prediction methods benefited from assessing a larger number of organisms. However, in line with earlier studies [34, 48], we found that increasing profile length beyond 500 organisms resulted in diminishing returns, supporting the observation that pathway discovery does not require many genomes.

**Figure 3.**
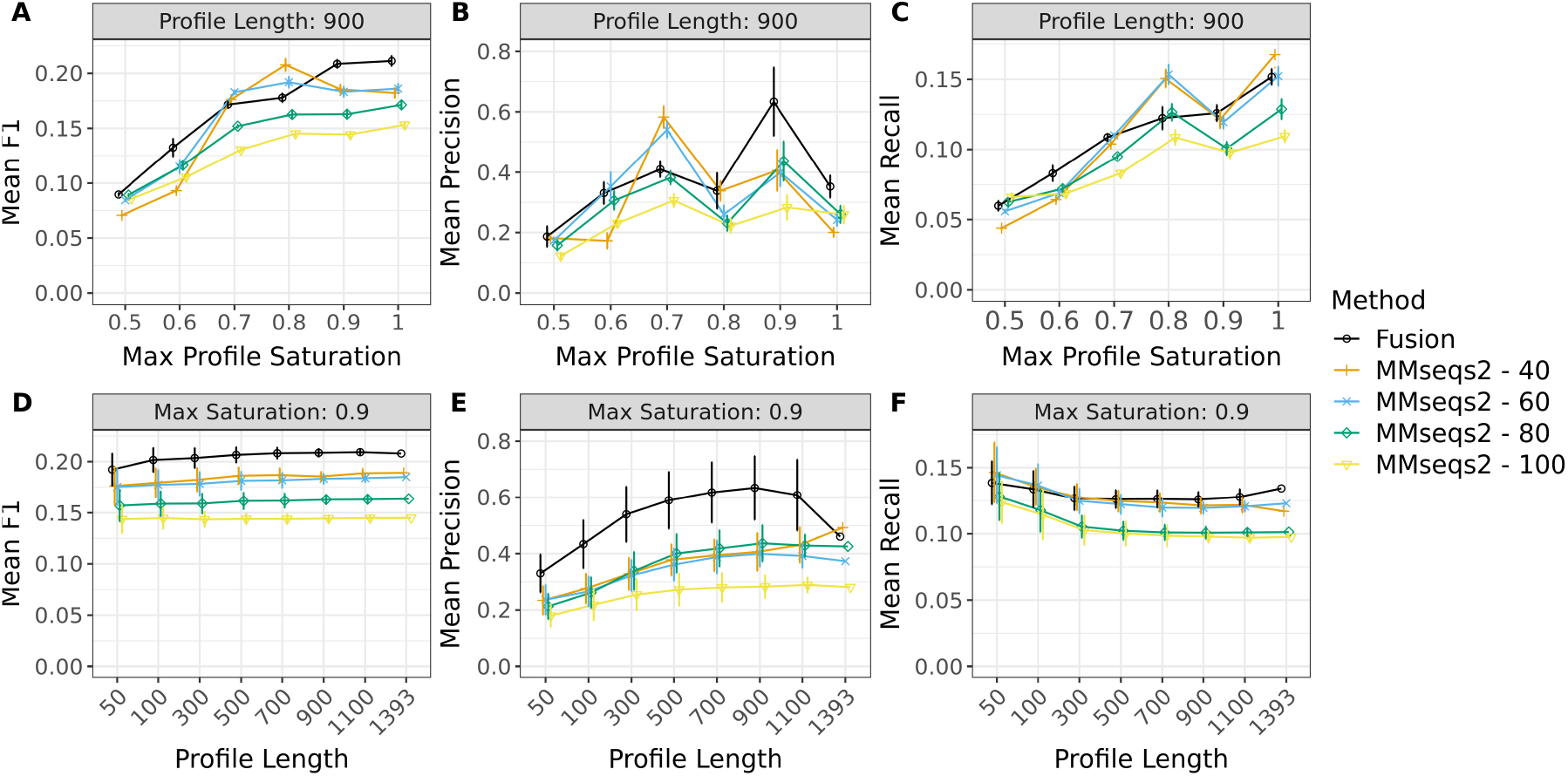
Predicting Protein Interactions at Varying Profile Size and Saturation Thresholds. We varied the length and the saturation of *fusion* (black lines) and MMseqs2 (colored lines, number indicates bit score of 40-100) profiles to identify the parameters for the best predictive performance. Error bars indicate standard deviation over 100 replicates; only one replicate was used to assess profile lengths of 1,393. In **(A-C)**, all values are shown using a profile length of 900 and explore the impact of saturation changes. In **(A)**, the mean *F*_1_ was highest for *fusion*, above 0.9 saturation; for MMseqs2, it was highest at lower bit scores (40, 60) and only at 0.8 saturation. Note that, at its highest, *fusion F*_1_ was better than that of MMseqs2. **(B)** and **(C)** show the mean precision and mean recall, respectively. For *fusion*, precision was maximized at saturation levels at or above 0.9 and recall was maximized by retaining all proteins. For MMseqs2, the best mean *F*_1_ (at a saturation level of 0.8) was achieved by improved recall. **(D-F)** compare the performance of profiles at different lengths, with a max profile saturation of 0.9. The mean *F*_1_ **(D)** of either method does not vary significantly above profile lengths of 500. **(E)** and **(F)** show that the maximum precision is achieved at a profile length of 900, with a maximum recall at length 1,393.

The saturation filter aimed to remove predicted false positive interactions of proteins that participate in (different) essential bacterial pathways. Ubiquitous proteins have high profile similarities even if they are not found in the same functional pathway. For example, transcription and glycolysis machinery, while found in nearly all bacteria in our data, are obviously not part of the same KEGG pathway. Moreover, essential functions identified throughout the bacterial landscape are likely to be extensively annotated and do not require further analysis [60]. Note that the saturation filter removes proteins from consideration, necessarily reducing a method’s ability to recall all possible interactions. A saturation filter of 0.9, i.e. removing proteins found in at least 90% of organisms, slightly decreased the *F*_1_ (Ref Fig. 3A), but greatly improved the precision (Ref Fig. 3B) of the prediction for both *fusion* and MMseqs2 profiles.

*Fusion* and MMseqs2 profiles achieved their highest precision with a max saturation of 0.9 and a length of 900. As such, we used these parameters for the rest of this work when making predictions.

### 3.2 Identifying interacting proteins is complicated by the accepted metrics and gold standard data

As shown above (Ref Fig. 3A),*fusion* profiles outperformed MMseqs2 profiles, achieving an overall higher *F*_1_ and precision; best, at saturation thresholds of 0.9 and 1 (black line higher than colored lines). Furthermore, *fusion* profiles achieved a higher recall than MMseqs2 profiles at the same levels of precision (Ref Fig. 4). For all profiles, however, there was a steep decrease in precision for higher recall values (Ref Fig. S1).

**Figure 4.**
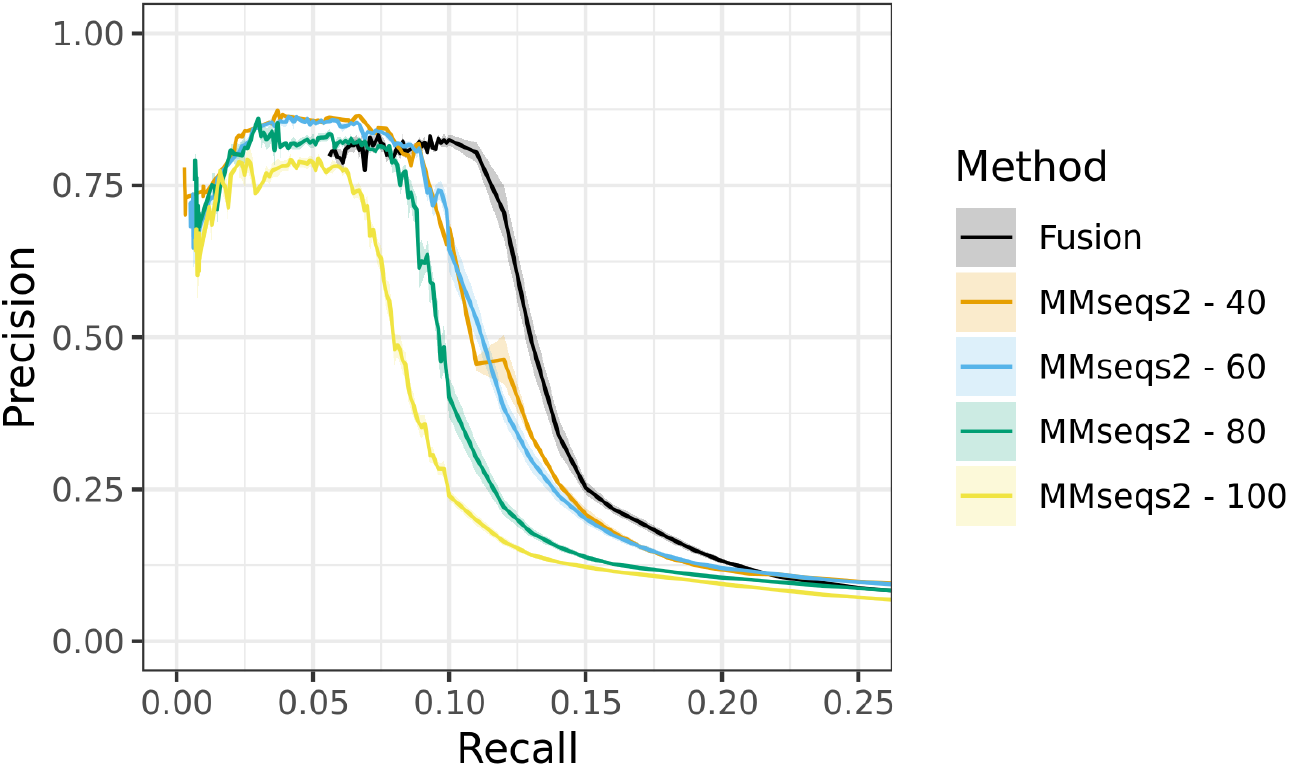
Comparing the Predictive Power of *fusion* and MMseqs2 profiles. At the same levels of precision (0.13-0.8), *fusion* profiles tend to recall more protein pairs from the same pathways than MMseqs2 profiles across multiple bit-score thresholds (40-100, colored lines).

In assessing Figure. 4, two issues are clear. (1) At lower recall values (*≤* 0.055), *fusion* profiles could not be directly compared to MMseqs. For a given protein, a *fusion* profile was generated by using all proteins from the same *fusion* cluster. Thus, profiles of proteins in the same cluster were necessarily the same, i.e. Jaccard Similarity = 1. Due to this limitation, 0.055 is the minimal recall for *fusion* profiles. Proteins of the same *fusion* cluster that are found in the same organism, are also likely paralogs. This does not suggest shared pathways and implies a further lowered precision even at these lower recall points. However, higher performance of *fusion* profiles across other areas of the Precision-Recall curve, lends confidence to our predictions.

(2) Both *fusion* and MMseqs2 achieved high prediction precision only in the areas of very low recall. Our definition of the ground truth explains this observation, i.e., the evaluation data set (Methods). We generated our Positives by labeling as interacting every protein pair within an organism module (Eqn. 1). This is untrue, as pathways tend to have distinct steps carried out by subsets of component proteins. As such, this approach leads to a drastic increase in the number of false pairwise interactions that each prediction method had to recapitulate. This inevitably biased our recall evaluations. Conversely, some proteins in distinct modules, e.g. M00001: Glycolysis and M00307: Pyruvate Oxidation, are intrinsically linked and separating these two modules as discrete processes is biologically minor. This inflates our Negatives with pairs of proteins that likely participate in the same or very similar biological processes but are in separate Modules.

To assess the effect of this bias in our evaluations, we compared the pairwise calculation of recall with a group-based measure of recall (Ref Fig. 5). Instead of focusing on the complete set of protein pairs, Group Recall is calculated by counting the fraction of proteins in a module that was correctly predicted to interact with one or more other proteins in the same module. Our results show that relative to Pair Recall, Group Recall is significantly higher and, as expected, increases with the number of proteins in a module. This latter observation indicates that at least one pair of interacting proteins is likely to be found in a larger module than in a smaller one.

**Figure 5.**
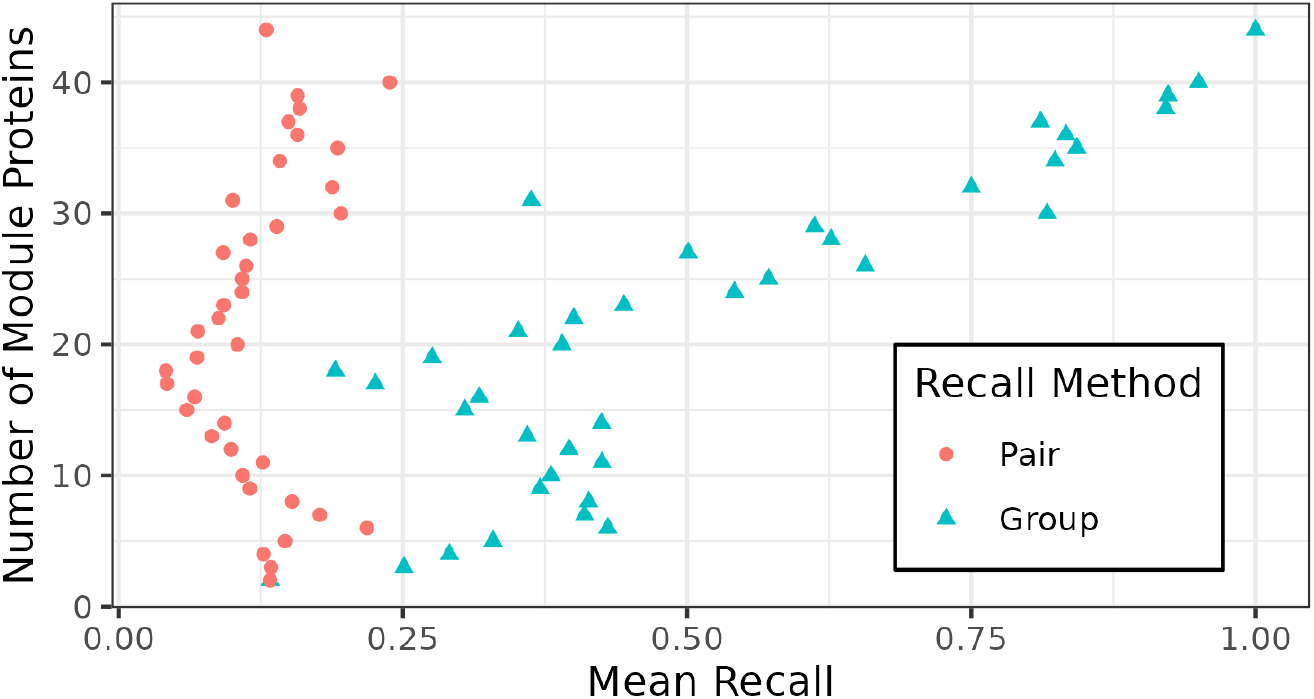
Pair and Group Recall Comparison. We compared the average Pair and Group Recalls for all modules of one size, i.e. number of proteins. The Pair Recall is much lower, especially for large module sizes (*≥* 20), due to the inflation of the number of putative Positive interactions.

Overall, the average Pair Recall for *fusion* predictions was 0.12, while Group Recall was 0.25. Though we used Pair Recall in our evaluations to facilitate comparing our results with prior studies, we propose that Group Recall is a more appropriate measure of evaluating pathway coverage.

### 3.3 Profiles of protein functional similarity describe pathways better than those based on sequence similarity

To understand why *fusion* profiles predict more functional relationships than MMseqs2 and whether this improvement is biased to a certain type of biological process, we gathered protein pairs in each module identified by both *fusion* and MMseqs2 (*fusion* bit-score = 40, i.e. the one with the highest *F*_1_ score, Ref Fig. 3). We then identified modules where *fusion* correctly predicted more pairs than MMseqs2, and calculated the significance of *fusion* predictions given the total number of possible protein pairs in each KEGG module (Bonferroni-Holm corrected for multiple comparisons [61]); we retained modules where the adjusted significance was *≤* 0.05. *Fusion* predicted significantly more interacting pairs than MMseqs2 in 52 different modules, including three modules where MMseqs2 predicted no interactions (Ref Tab. S2). We grouped these modules into their KEGG-assigned classes (Ref Tab. 1). The largest difference (fold change) between predictions was for pairs of proteins representing functions in Glycan metabolism. We repeated this analysis for modules where MMseqs2 correctly predicted more pairs than *fusion* and counted 19 modules, mostly from the Amino Acid (6 modules) and Energy (5 modules) metabolism pathways.

**Table 1:** Module classes predicted by *fusion* and MMseqs2 profiles. We summarized modules where *fusion* profiles predicted significantly more protein pairs than MMseqs2 by their module class. The number of protein pairs identified by each method is shown along with the fold change increase for *fusion* over MMseqs2. The largest fold change in performance for *fusion* profiles over MMseqs2 was for Glycan Metabolism and Signature-class modules. Signature modules represent pathways unique to a specific subset of organisms that are not grouped with the larger, more general classes - specifically, Nitrate and Sulfate-sulfur assimilation. Fusion had a higher recall in these modules than MMseqs2, uniquely predicting more protein pairs in environmental processes.

| Module Class | Total Possible Pairs | MMseqs2 | Fusion | Fold Change |
| --- | --- | --- | --- | --- |
| Glycan metabolism | 17173 | 81 | 2565 | 31.7 |
| Signature modules | 8524 | 31 | 713 | 23 |
| Biosynthesis of other secondary metabolites | 150 | 1 | 21 | 21 |
| Gene set | 514 | 3 | 42 | 14 |
| Xenobiotics biodegradation | 10851 | 132 | 602 | 4.56 |
| Energy metabolism | 63926 | 1881 | 5589 | 2.97 |
| Carbohydrate metabolism | 23870 | 695 | 1632 | 2.35 |
| Lipid metabolism | 32487 | 1781 | 3466 | 1.95 |
| Metabolism of cofactors and vitamins | 51359 | 5370 | 9832 | 1.83 |
| Amino acid metabolism | 259565 | 28919 | 46140 | 1.6 |
| Biosynthesis of terpenoids and polyketides | 2219 | 369 | 371 | 1.01 |

We then asked if there were any shared quantitative characteristics of the modules that could explain the difference in method performance. Using a Wilcoxon signed-rank test, we compared (1) the number of possible protein pairs in each module; (2) the number of unique enzymatic protein functions at third and fourth EC levels; (3) the Shannon diversity of functions at third and fourth EC levels; (4) the number of organisms in which a module is found; (5) the median base-pair distance between module genes along the corresponding chromosomes; (6) the median *fusion* cluster size of participating proteins; (7) the number of KEGG orthologs in the module; (8) the number of module steps or sequential enzymatic reactions that comprise a module, and (9) the number of non-orthologous replacements (NORs), i.e. the number of unique KOs that catalyze a step in the module.

The Shannon diversity *H*(*X*) of EC numbers (*x*) in each module was calculated 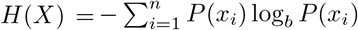, with *P* (*x*_*i*_) representing the probability of the *i* -th EC number in the module. The p-value was corrected for multiple comparisons using the Bonferroni-Holm correction [61]; a significance threshold of 0.05 was applied to the adjusted p-value.

We found no significant difference in any of the tested attributes (Ref Fig. S2). That is, we could not identify a significant pattern in the biological class or characteristic numeric features of the predicted modules that would benefit *fusion*-based predictions.

We further explored the possible mechanistic differences in the structure of *fusion* and MMseqs2 profiles. We assessed the 62,536 profiles of proteins participating in the modules where *fusion* predicted more relationships than MMseqs2. Of these, two-thirds (41,196 proteins) had lower saturation *fusion* than their equivalent MMseqs2 profiles. The median saturation for *fusion* profiles was 0.69 and 0.74 for MMseqs2 profiles. However, this observation was not specific to only these proteins. Over our complete set of 306,377 proteins, the median saturation of all *fusion* and MMseqs2 profiles was 0.65 and 0.77, respectively. We reason that this results from *fusion* more precisely identifying functionally-similar rather than sequence-similar proteins across organisms.

To support our reasoning, we calculated the Rand Index (Eqn. 3) of *fusion* and MMseqs2 profile groups against five different measures of functional similarity. As expected from our earlier work [41, 42], *Fusion* met or exceeded MMseqs2’s ability to correctly label proteins across all measures, namely; sharing the same third- (1) or fourth- (2) level EC number (505 proteins) (3) participating in the same KEGG module (306,377 proteins); (4) assigned the same KO label (306,377 proteins), and (5) interchangeable in a module as non-orthologous replacements (NORs, 2,200 proteins) (Ref Tab. 2).

**Table 2:** Rand Index Comparison for Functional Annotations. The Rand Index, i.e. percent of protein pairs with the same functional annotation (EC, Module ID, KO, or NOR) of protein groups labeled with *fusion* and MMseqs2. *Fusion* more accurately identified proteins as sharing the function label across all categories.

|  | EC4 | EC3 | Module ID | KO | NOR |
| --- | --- | --- | --- | --- | --- |
| **Fusion** | 67.7% | 98.9% | 71.1% | 67.3% | 1.64% |
| **MMseqs** | 55.4% | 93.8% | 62.4% | 42.1% | 1.60% |

The performance improvement, yielded by *fusion*’s precise functional grouping of proteins, was not replicated by increasing the bit-score threshold of MMseqs2 profiles. In fact, a bit-score threshold of 100, the most stringent measure used in our analysis, had the lowest overall performance. This observation is not unexpected, as sequence dissimilar proteins may share the same function [62]. In fact, the median pairwise identity of proteins in one *fusion* cluster was 34.7%, and the median identity of MMseq2-aligned protein pairs at bit-score thresholds of 40, 60, 80 and 100 were 32.9%, 34.6%, 36.4%, and 38.2% respectively (Ref Fig. S3). That is, prediction precision is not strictly a function of sequence identity.

Note that both *fusion* and MMseqs2 profiles recall relatively few of our gold standard interacting proteins (Ref Fig.3). These results are not unexpected, as the precision of phylogenetic profiles is known to decrease with higher recall [53]. The presence of homologs born from gene duplication or horizontal gene transfer (HGT), as well as multi-functional proteins that participate in multiple metabolic pathways, interferes with some of the underlying assumptions of our profiles. For example, sequence similarity-based profiles will, by definition, be fooled by paralogs that share the same phylogenetic profile but participate in distinct pathways.

We further looked for a set of proteins that exemplified our reasoning in favoring functional profiles over sequence similarity-based ones. We found an example when evaluating the profiles of AFY38555, a tocopherol cyclase protein in *Enugrolinea bermudensis PCC 7376*. Tocopherol biosynthesis is one of 18 KEGG modules for which *fusion* identified interacting protein pairs that went undetected by MMseqs2. As expected, *fusion* profiles for tocopherol biosynthesis proteins were more precise, containing fewer members than their MMseqs2 equivalents. The *fusion* cluster for AFY38555 included 32 elements corresponding to labeled tocopherol cyclases, putative tocopherol cyclases, and hypothetical proteins found exclusively in Cyanobacteria. The MMseqs2 AFY38555 matches included CDR30235, in addition to the proteins identified by *fusion*. CDR30325 is a toco-pherol cyclase from the Mycoplasmatotoa, *Acholeplasma oculi*. Thus, although MMseqs2 identified a protein of an apparently correct function, the MMseqs2 profile had lower profile similarity with other tocopherol cyclase proteins due to its function being specific in a separate clade. We conclude that *fusion’s* ability to identify functionally similar proteins with greater precision than MMseqs2 contributes to more accurate profiles and improved functional inference.

### 3.4 Fusion similarity-based networks describe complete molecular pathways

We constructed and clustered both *fusion* and MMseqs2 protein profile similarity networks (Methods) to generate multi-protein interaction predictions of potential pathways. Predicted pathways ranged in size from 2–98 proteins, where most 2-protein clusters constituted either proteins of the same putative function or sub-units of larger multimeric protein complexes. Of the 22,756 generated 2-protein pathway predictions, 76% were assigned to the same *fusion* cluster. A review of the larger clusters revealed several known interactions between proteins, most commonly in multiple amino acid biosynthesis and photosystem pathways. We evaluated all clusters by calculating the maximal Jaccard similarity between a predicted cluster and each possible KEGG pathway. *Fusion*-based pathway predictions had a significantly higher maximal Jaccard similarity with known KEGG pathways than clustered MMseqs2 profiles (*fusion* median = 0.47, sd = 0.21; MMseqs2 median 0.37, sd = 0.22; *p − val ≤* 2.2*x*10^*−*16^, Mann-Whitney Utest). Furthermore, we observed that, while *fusion* predictions were better than sequence-based ones across all modules, those comprised of 2–5 proteins were most difficult for MMseqs2 profiles (Ref Fig. 6). We thus conclude that *fusion* clusters, which identify functionally similar proteins, improve bacterial pathway prediction over MMseqs2, our sequence similarity-based comparison method.

**Figure 6.**
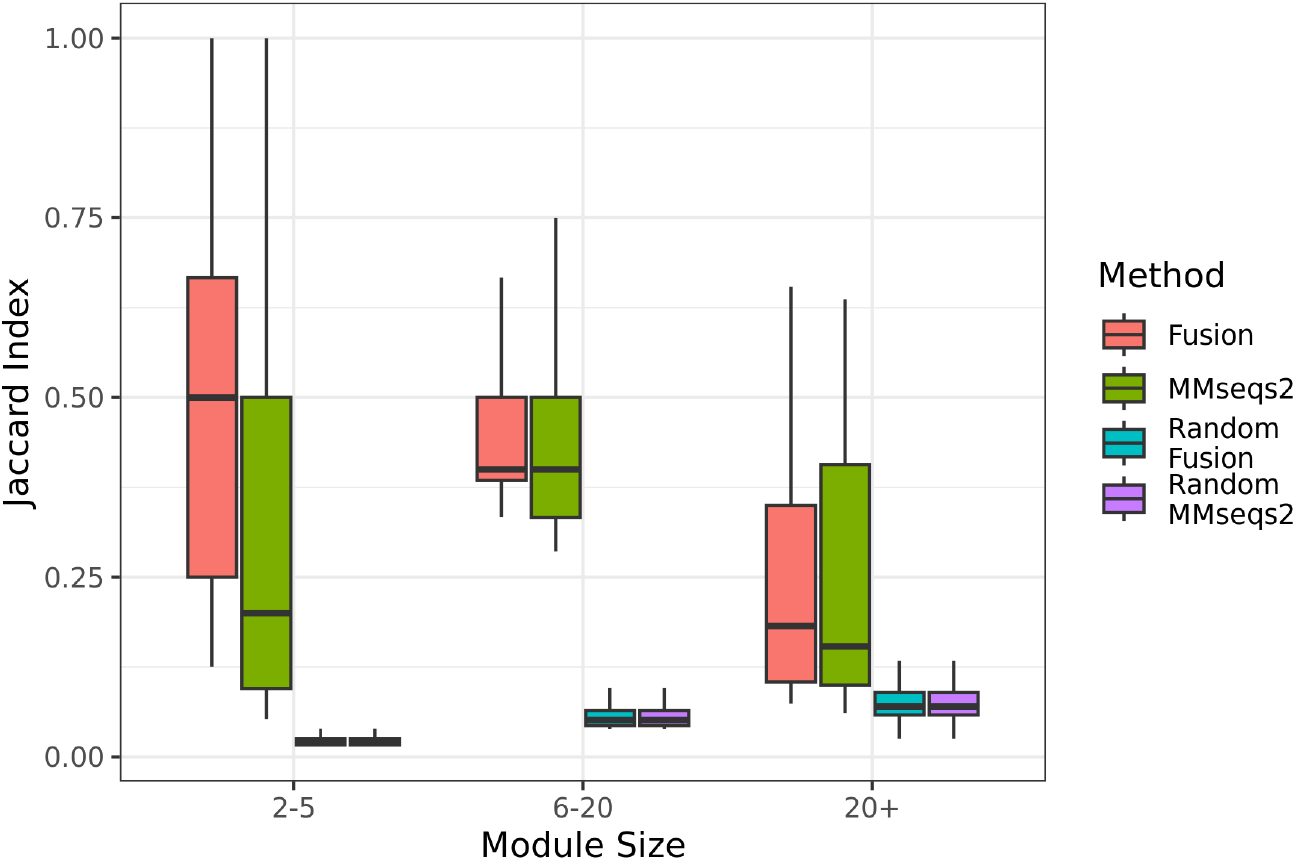
Maximal Jaccard Similarity of *fusion* and MMseq2 -labeled pathway clusters. Clusters of proteins based on *fusion* profile similarities have a high similarity (Y-axis) to KEGG modules in bacterial organisms. The X-axis separates modules by the total number of proteins in KEGG’s organism-specific module. The color of each boxplot indicates the method used to generate clusters and their randomized comparisons; red (fusion), green (MMseqs2), blue (randomized *fusion*), and purple (randomized MMseqs2). A total of 62,378 KEGG modules were considered, and the top 10,000 Jaccard similarity values are shown. A full comparison is shown in (Ref Fig. S4)

The previous results demonstrate *fusion’s* advantage when predicting pathway components from proteins of the same organism. To validate the power of our approach in finding emergent pathways, we similarly clustered a larger network of all proteins in our set. We compared the resulting clusters against all KEGG pathways, regardless of the organism of origin. While somewhat less so, *fusion* still outperformed MMseqs2; median Jaccard scores of 0.21 (sd = 0.17) and 0.15 (sd = 0.15), respectively.

### 3.5 Metagenomic profiles reveals unique putative functional relationships between proteins

We extended our use of functional co-occurrence to identify emergent pathways in metagenomic data. Co-occurrence networks have previously been used in metagenome studies to predict novel interactions between organisms [63–68]. The co-occurrence of genetic elements, such as antibiotic resistance genes or phages, has also been applied to metagenomes [69–72]. Here, we used *fusion* functional annotations to pinpoint co-occurring protein functions in marine microbiome samples directly from short-read data. By additionally using taxonomic annotations, we split up the task of looking for emergent functionality into multiple steps: (1) identify pathways that we expect to occur and be present based from taxonomic inference; (2) identify pathways present but not attributable to any annotated taxa; (3) predict putative pathways made up of known functions but not found in any single known species, and (4) predict putative pathways between proteins of unknown function. In this step-by-step manner, we assess both the validity and usefulness of our approach for understanding community pathways. We also propose a means for inferring the degree of functional interconnectedness in microbial communities without the strict need for functional annotation.

We used Kaiju to generate taxonomic annotations for marine microbiome samples from the bioGEOTRACES project (PRJNA385854, Methods) [73]. The majority of reads in each sample, 66%-93%, were unable to be annotated, as is often the case [69, 74–76]. Of the annotated reads, Proteobacteria, Bacteroidota, and Cyanobacteria were the most prevalent phyla. We limited the inference of functions to species inferred from at least 1% of annotated reads to give some certainty in our detection of present functions. We cross-referenced the detected bacterial strains against KEGG, and inferred the corresponding KEGG modules and component enzymatic proteins. Of 480 samples, eight contained no species (with a corresponding KEGG entry) representing *≥* 1% of the total diversity. The lack of dominant taxa in these eight samples highlights the benefit of direct functional annotations. For the remaining 472 samples, a median of 58 different modules and 218 unique EC numbers were extracted for each sample. The total number of unique modules and EC numbers identified across all samples were 188 and 674, respectively. These values quantify the functional potential of the metagenome samples able to be attained from taxonomic annotation and subsequent inference of function.

For our taxonomically independent, function-focused annotation, we used *mi-faser* [43]. We identified a median of 267 EC numbers per sample. We note that this number of enzymatic functions detected from the direct annotation of reads exceeds the Kaiju-estimated number (218) reported above. This finding is unsurprising given the large fraction of reads not assigned to any specific taxa. Furthermore, for each sample, the two methods overlapped by only a median 69 EC numbers; a median of 198 and 144 EC numbers were identified exclusively by *mi-faser* or Kaiju, respectively. In light of the previously reported *mi-faser* performance — high precision (0.94) and moderate recall (0.52) — we expect that both *mi-faser* and Kaiju predictions may be correct for a significant fraction of annotations.

We explored the number of complete modules, determined by the presence of all of the constituent EC numbers predicted by *mi-faser* [43] in a given sample, and identified 2,321 complete modules across all samples. We then repeated this evaluation for ‘unexpected’ modules, or KEGG modules belonging to organisms not annotated to be present by Kaiju, yielding 2,139 complete modules despite the expectation that the module would be absent given the present taxa. As a comparison for our pathway detection, we repeated these module-detection steps with a random sample of detected EC numbers and only recovered 84 and 590 complete expected and unexpected modules, respectively. Our results suggest that the direct annotation approach yielded the correct enzymatic building blocks to construct bacterial pathways. With these building blocks identified, we proceeded with reconstructing pathways with our profiling approach.

Our emergent pathway reconstruction approach uses the same principles as the profile analysis described above. We created metagenomic profiles, representing the occurrence of each enzymatic function in each sample, and clustered these profiles by Jaccard similarity. In this case, we did not apply a saturation filter to the profiles as we did with our initial evaluation. The purpose of the saturation filter was to remove from consideration proteins ubiquitous to the full diversity of organisms in our data set — a notion not useful for analysis of our environmentally homogeneous marine metagenome samples. That is, in this application, we instead deliberately sought out pathways core to our set of marine metagenomes, instead of pathways unique to a subset. In total, we generated 12 protein clusters based on co-occurrence patterns across 472 samples. Note that we expected that having fewer metagenomic samples relative to our *fusion* profiles (shorter length) would not hamper our inference, given the marginal loss in performance shown in our evaluation of profile lengths. We assessed the resulting clusters to find 11 expected and 12 unexpected modules, respectively.

Having been successful in recapitulating pathways exclusive to individual organisms, we aimed to detect pathways between organisms, i.e. *emergent pathways*. We repeated our metagenomic profile clustering with the additional step of filtering edges to retain pairs of proteins not known to co-occur in the proteome of any taxa identified by Kaiju (Ref Tab. S3). We produced 11 clusters whose proteins overlapped with 20 additional complete modules not found in any of the Kaiju-identified species according to their KEGG entries, including: four Energy metabolism, four carbohydrate metabolism, three Xenobiotics biodegradation, three Metabolism of cofactors and vitamins, three Amino acid metabolism, and one each of Nucleotide metabolism, Lipid metabolism, and Biosynthesis of terpenoids and polyketides class modules. This analyses demonstrates the ability of our method to recover emergent biochemical pathways based on the co-occurrence of function.

Finally, we examined our clusters for emergent interactions between unknown functions not found in any KEGG modules, as per our fourth pathway detection step described above. The purpose of this final evaluation was to establish a basis for which currently unknown biochemical pathways between unknown functions could be prioritized based on their ease of discovery by co-occurrence analysis. We reviewed the generated clusters to find any previously unseen interactions that could be supported by our understanding of biochemical pathways in bacteria. Among the emergent pathways was a predicted interaction between dihydroorotate dehydrogenase (DHODH, EC: 1.3.5.2) and nucleoside-2’,3’-cyclic-phosphate 3’-nucleotidohydrolase (CNP, EC: 3.1.4.16), both of which participate in pyrimidine metabolism. In gram-negative bacteria, CNP is localized to the periplasm, where it assists in the conversion of 2’,3’-cyclic-nucleotides into 3’-nucleotides [77, 78]. This functionality is necessary as bacteria cannot transport 2’,3’-cyclic-nucleotides into the cytoplasm, and thus require their conversion to 3’-nucleotides, followed by subsequent steps for intake. Putatively, CNP functions as a scavenger of exogenous nucleotides, and cloning of the CNP gene in *Yersinia entercoliticia* has enabled its subsistence on 2’,3’-cAMP as a sole source of carbon and energy [79, 80]. Consistent, across multiple metagenomes, co-occurrence of CNP with DHODH, a critical enzyme for *de-novo* pyrimidine biosynthesis, suggests a similar process, whether in a single undetected bacterium or shared between multiple ones, could be occurring in the marine microbiome. Earlier studies have found pyrimidine and purine metabolism to be among the most abundant pathways in marine bacterial communities [81]. Thus, the production of pyrimidine and its further conversion into extracytoplasmic cyclic dinucleotides may be responsible for availing an energy source to CNP containing bacteria.

In addition to the biochemical pathways intrinsic to individual microbes, microbiomes host various shared/emergent interactions between bacteria. Detection of these emergent pathways allows for better and higher resolution annotation of the microbiome’s functional potential. Here we propose a method to predict emergent pathways, generating testable hypotheses of essential protein interactions in different microbial environments. We emphasize that unique to our study is the generation of these hypotheses incorporating proteins of both known and unknown function. In agreement with a prior study of the marine metagenomes [82], we clarify the mechanisms regulating microbial communities by differentiating the discernible functions in a sample from taxonomy. Understanding microbiome functional mechanisms is crucial for assessing how microbiomes change in response to environmental conditions [83, 84], improving mathematical models of microbial systems [85, 86], and illuminating microbiome functional redundancies [87–89].

Beyond identifying possible emergent pathways containing functionally annotated proteins, we identified 46 clusters/putative pathways, of functionally unannotated proteins. We thus provide testable hypotheses for the experimental targeting of the yet to be identified functional potential of microbiomes. Considering the rate of growth of metagenome sequences collected from a variety of locations and conditions, we suggest that our functional analyses that forgo taxonomy, do not require genome assembly, and even make use of yet unknown functions will produce valuable and previously inaccessible, insights into the functionality of microbial communities.

## 4 Conclusions

The expanding reservoir of metagenomic sequences promises to yield an unprecedented windfall for microbial discovery. Extracting biological information from these sequences remains a challenge even as data availability skyrockets [90]. Here, we used *fusion*, a protein functional annotation method that does not rely on reference databases, to predict molecular pathways based on function co-occurrence. *Fusion* profiles, describing microbe-specific proteins of shared function, identified more functional interactions than a comparable sequence-homology-based method. We further applied our approach to metagenome samples, identifying a putative emergent energy metabolism pathway and predicting additional cross-organism pathways that have yet to be validated. These emergent pathways allow for the subsequent characterization of microbiomes by their functional potential, an essential step toward building a more complete understanding of microbial communities.

## Code and Data Availability

The code used to generate profiles and subsequent analyses in this study is available on GitHub; https://github.com/FriedbergLab/PAMFUN

The datasets generated and analyzed in the study, including; *fusion* and MMseqs2 profiles, KEGG reference, and pathway predictions, are available in the following Figshare repositories; https://doi.org/10.6084/m9.figshare.25066478.v1, https://doi.org/10.6084/m9.figshare.25066163.v1, https://doi.org/10.6084/m9.figshare.25025231.v1.

