## Supplementary figures and images for "Assembling bacterial puzzles: piecing together functions into microbial pathways"

### Supplemental Figure 1

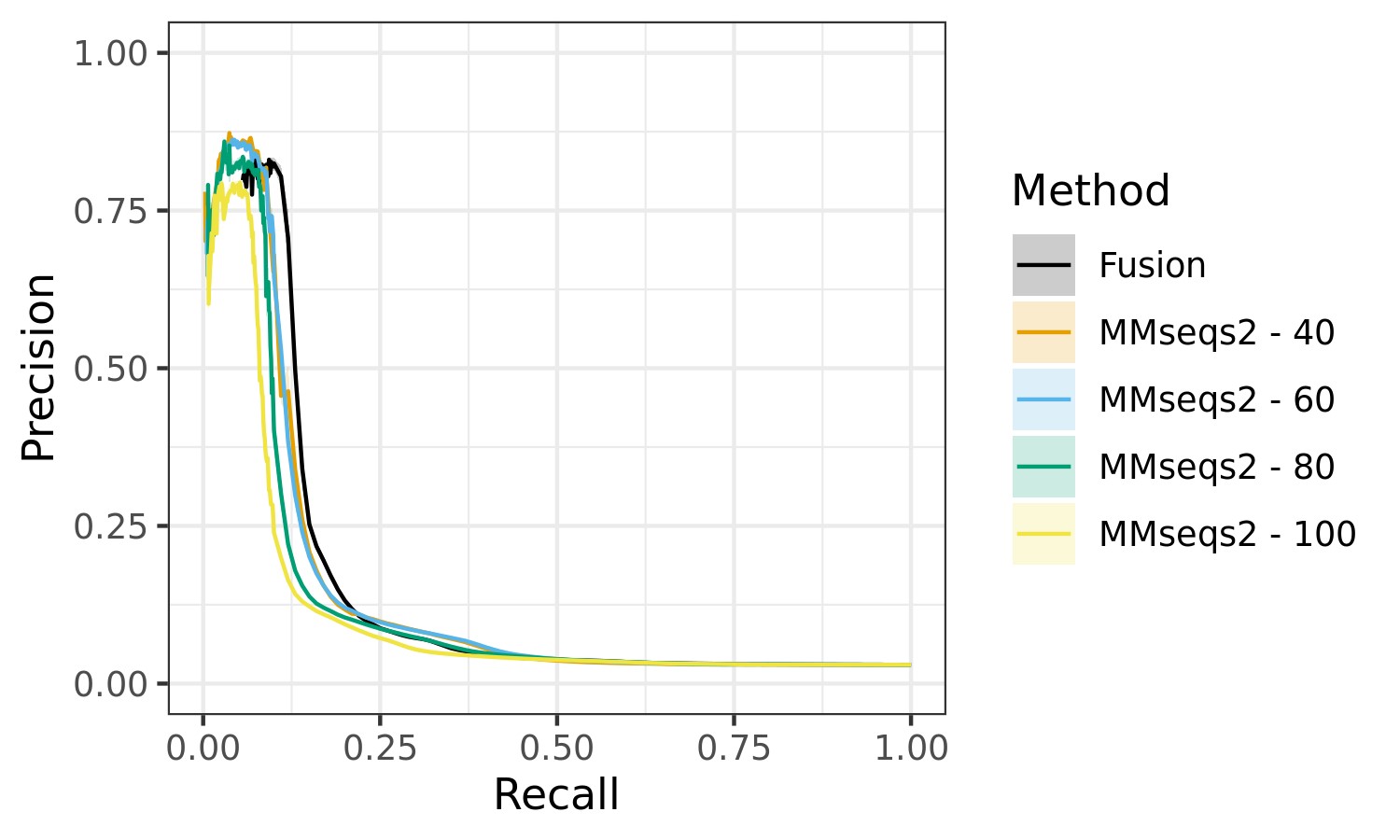

### Supplemental Figure 2

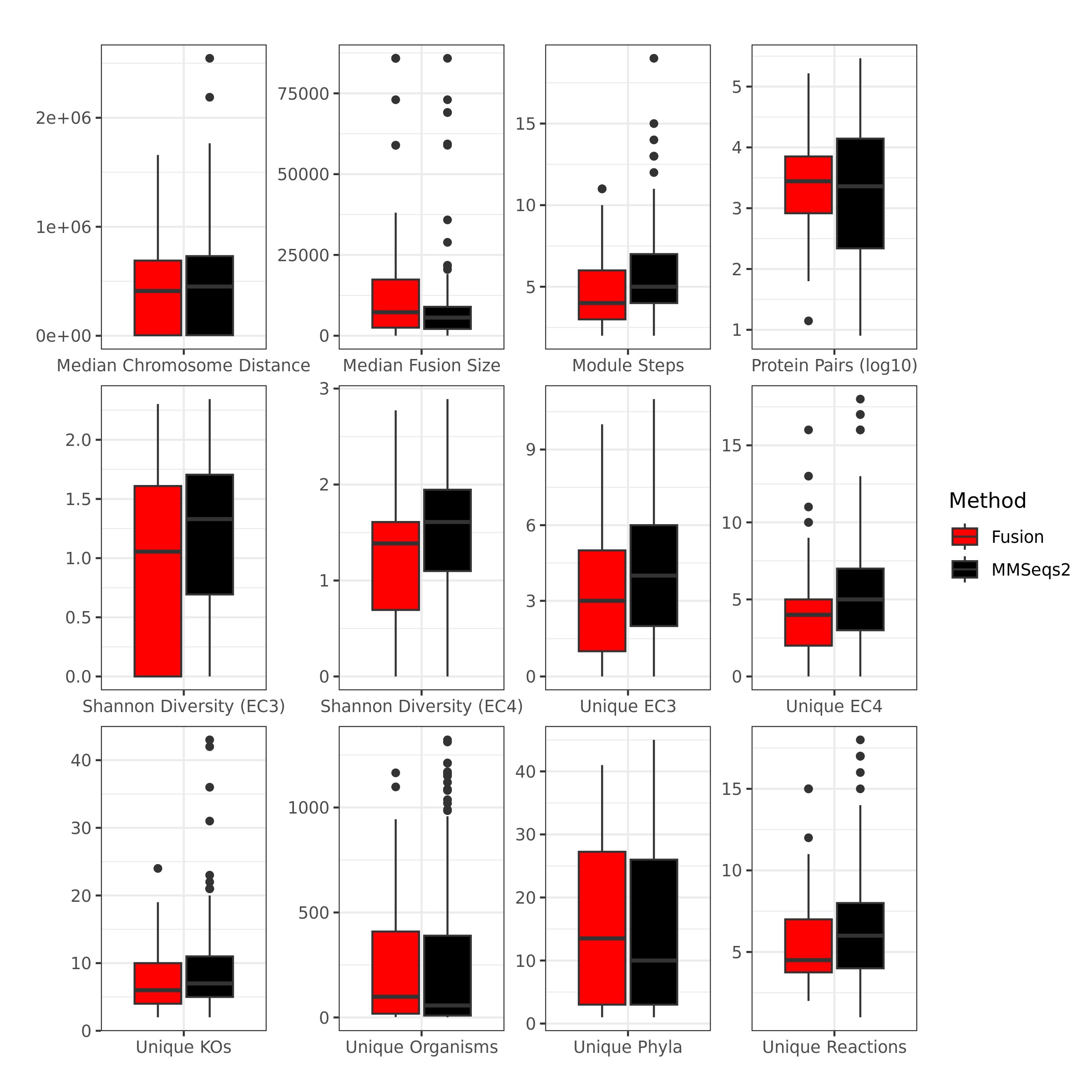

### Supplemental Figure 3

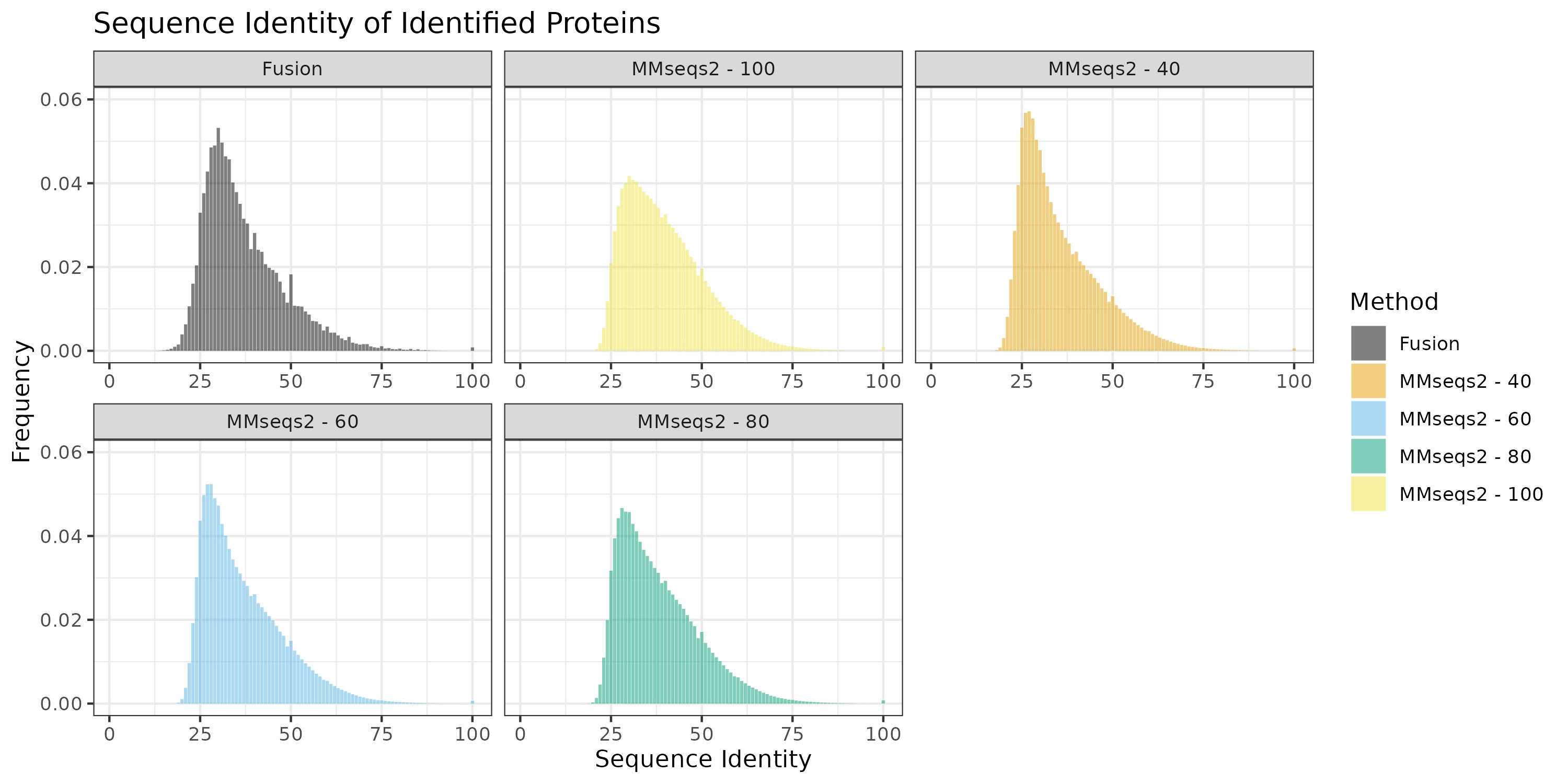

### Supplemental Figure 4

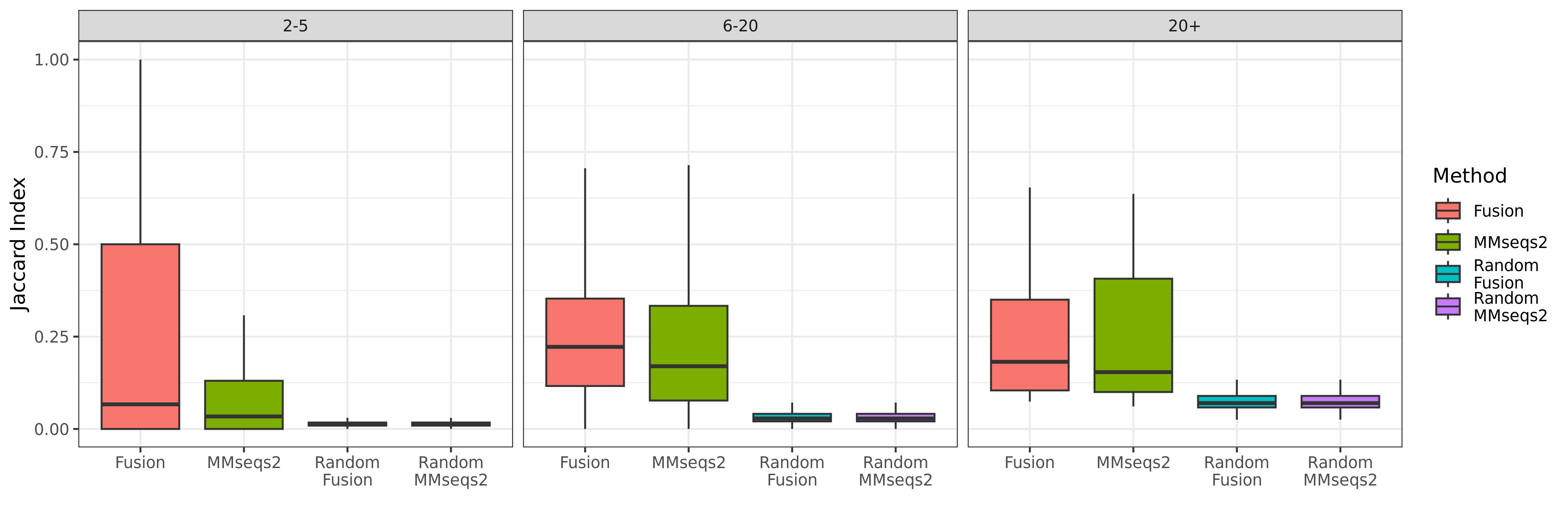
